# Is coding a relevant metaphor for the brain?

**DOI:** 10.1101/168237

**Authors:** Romain Brette

**Affiliations:** Sorbonne Universités, UPMC Univ Paris 06, INSERM, CNRS, Institut de la Vision, 17 rue Moreau, 75012 Paris, France

**Keywords:** neural coding, information, perception, sensorimotor, action

## Abstract

I argue that the popular neural coding metaphor is often misleading. First, the “neural code” often spans both the experimental apparatus and the brain. Second, a neural code is information only by reference to something with a known meaning, which is not the kind of information relevant for a perceptual system. Third, the causal structure of neural codes (linear, atemporal) is incongruent with the causal structure of the brain (circular, dynamic). I conclude that a causal description of the brain cannot be based on neural codes, because spikes are more like actions than hieroglyphs.

**Long abstract:** “Neural coding” is a popular metaphor in neuroscience, where objective properties of the world are communicated to the brain in the form of spikes. Here I argue that this metaphor is often inappropriate and misleading. First, when neurons are said to encode experimental parameters, the neural code depends on experimental details that are not carried by the coding variable. Thus, the representational power of neural codes is much more limited than generally implied. Second, neural codes carry information only by reference to things with known meaning. In contrast, perceptual systems must build information from relations between sensory signals and actions, forming a structured internal model. Neural codes are inadequate for this purpose because they are unstructured. Third, coding variables are observables tied to the temporality of experiments, while spikes are timed actions that mediate coupling in a distributed dynamical system. The coding metaphor tries to fit the dynamic, circular and distributed causal structure of the brain into a linear chain of transformations between observables, but the two causal structures are incongruent. I conclude that the neural coding metaphor cannot provide a basis for theories of brain function, because it is incompatible with both the causal structure of the brain and the informational requirements of cognition.

## 1. Introduction

A pervasive paradigm in neuroscience is the concept of neural coding (deCharms and Zador, 2000): the query “neural coding” on Google Scholar retrieves about 15,000 papers in the last ten years. Neural coding is a communication metaphor. An example is the Morse code (Fig. 1A), which was used to transmit texts over telegraph lines: each letter is mapped to a binary sequence (dots and dashes). In analogy, visual signals are encoded into the spike trains of retinal ganglion cells (Fig. 1B). Both the Morse code and the retinal code relate to a communication problem: to communicate text messages over telegraph lines, or to communicate visual signals from the eye to the brain. This problem has been formalized by communication theory (Shannon, 1948), also called information theory, a popular tool in neuroscience (Rieke et al., 1999).

**Figure 1.**
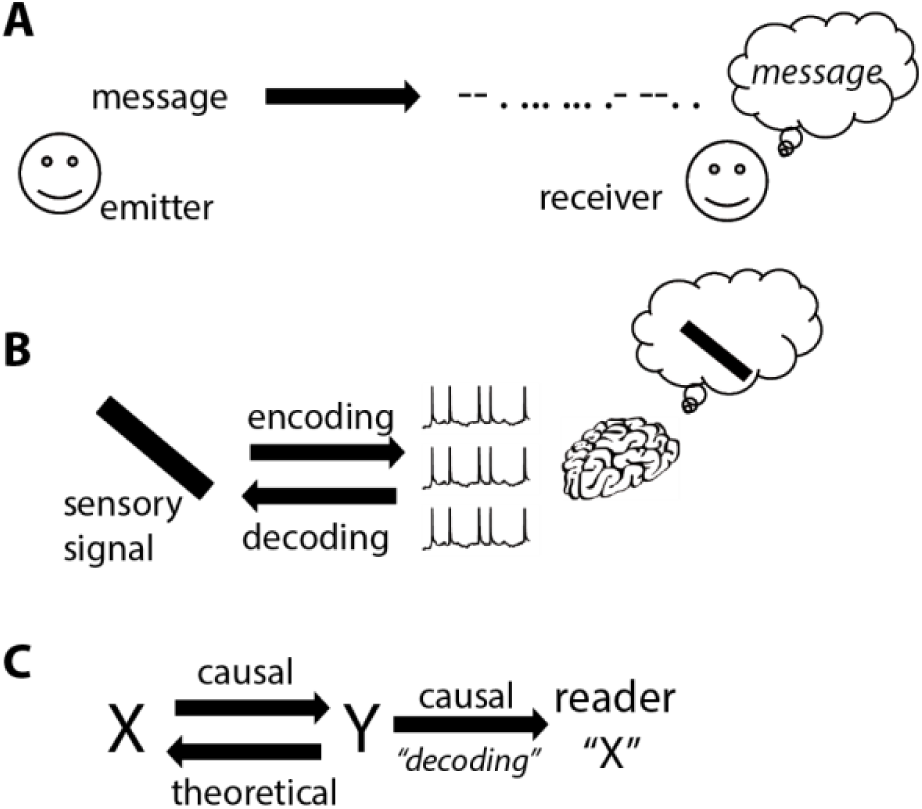
The coding metaphor. A, An emitter transmits a message to a receiver, in an altered form named “code” (here Morse code). The receiver knows the correspondence and can reconstruct (“decode”) the original message. B, In analogy, visual signals are encoded in the spike trains of the optic nerve. The rest of the visual system treats these spike trains as visual information. C. Implicit structure of the neural coding metaphor (“Y encodes X”): there is a correspondence between X and Y; encoding refers to a causal mechanism from X to Y, while decoding is a theoretical inverse mapping; Y causes changes in the reader (often improperly called “decoding”), and represents X in some sense.

The neural coding metaphor has shaped neuroscience thinking for more than five decades. Barlow (1961) used the metaphor extensively in his work on sensory neurons, although he warned to *“not regard these ideas as moulds into which all experimental facts must be forced”*. In a seminal review entitled “Neural coding”, Perkel and Bullock (1968) depicted *“the nervous system [as] a communication machine”* and already recognized the *“widespread use of “code”in neuroscience”*. An illustration of hieroglyphs figures prominently at the top of the technical appendix. Around the same time entire books were devoted to “sensory coding” (Somjen, 1972; Uttal, 1973).

As the linguists Lakoff and Johnson (1980) have argued, the metaphors that pervade our language are not neutral; on the contrary, they form the architecture of our conceptual system. What are the concepts carried by the neural coding metaphor, which makes it a possibly relevant metaphor for the activity of the retina? There are three key properties (Fig. 1C), which are all used in Perkel and Bullock’s review:

1) The technical sense of a code is a correspondence between two domains, e.g. visual signals and spike trains. We call this relation a code to mean that spike trains specify the visual signals, as in a cipher: one can theoretically reconstruct the original message (visual signals) from the encoded message (spike trains) with some accuracy, a process called *decoding*. Information theory focuses on statistical aspects of this correspondence (Shannon, 1948). It is in this sense that neurons in the primary visual cortex encode the orientation of bars in their firing rate, neurons in the auditory brainstem encode the spatial position of sounds (Ashida and Carr, 2011), and neurons in the hippocampus encode the animal’s location (Moser et al., 2008).

2) Yet, not all cases of correlations in nature are considered instances of coding. Climate scientists, for example, rarely ask how rain encodes atmospheric pressure. Another key element of the coding metaphor is that the spike trains are considered messages for a reader, the brain, about the original message: this is the representational sense of the metaphor. Perkel and Bullock call the reader’s activity *“interpretation of the encoded information”*. In his book on sensory coding, Somjen (1972) writes: “*Information that has been coded must at some point be decoded also; One suspects, then, that somewhere within the nervous system there is another interface […] where ‘code’ becomes ‘image.’”*. Similar statements abound in modern neuroscience literature: *“A stimulus activates a population of neurons in various areas of the brain. To guide behavior, the brain must correctly decode this population response and extract the sensory information as reliably as possible.”* (Jazayeri and Movshon, 2006).

3) Finally, we would not say that visual signals encode retinal spike trains, even though this would comply with the technical sense. The reason is the communication metaphor implicitly assumes a causal relation between the original message and the encoded message; here, spike trains result from visual signals by a causal process (transduction). Similarly, to be a representation for a reader, the neural code must at least have a causal effect on the reader. This causal structure is implicit in Perkel and Bullock’s definition of neural coding: *“the transformations of information in the nervous system, from receptors through internuncials to motor neurons to effectors”*.

These three elements (correspondence, representation, causality) constitute the conceptual scaffold of the neural coding metaphor. It could be argued that most technical work on neural coding only uses the first technical sense (correspondence), where the word “code” is used as a synonym for “correlate”. The use of the metaphor would thus only amount to an inappropriate but innocuous choice of words. But what is the scope of neural codes if they have no causal powers? In his famous critique of Skinner’s behaviorism, Chomsky (1959) summarizes the problem with the improper use of metaphors: *“[Skinner] utilizes the experimental results as evidence for the scientific character of his system of behavior, and analogic guesses (formulated in terms of a metaphoric extension of the technical vocabulary of the laboratory) as evidence for its scope*”. The goal of this article is to show that this quote fully applies to the neural coding metaphor, as used in various theories of brain function.

The general argument is as follows. Scientific claims based on neural coding rely on the representational sense or at least on the causal sense of the metaphor. But none of these two senses is implied by the technical sense (correspondence). When we examine the representational power of neural codes (part 1), we realize that coding variables are shown to correlate with stimulus properties but the code depends on the experimental context (stimulus properties, protocol, etc). Therefore neural codes do not provide context-free symbols. But context cannot be provided by extending the code to represent a larger set of properties, because context is what defines properties (e.g. the orientation *of a bar*). Thus, neural codes have little representational power. The fundamental reason (part 2) is that the coding metaphor conveys an inappropriate concept of information. Neural codes carry information by reference to things with known meaning. In contrast, perceptual systems have no other option than to build information from relations between sensory signals and actions, forming a structured internal model. But neural codes have no or little structure, so they are inadequate for this purpose. Finally (part 3), the neural coding metaphor tries to fit the causal structure of the brain (dynamic, circular, distributed) into the causal structure of neural codes (atemporal, linear), substituting the arbitrary temporality of algorithms for the temporality of the underlying physical system. The two causal structures are incongruent. Without denying the usefulness of information theory as a technical tool, I conclude that the neural coding metaphor cannot constitute a general basis for theories of brain function because it is disconnected from the causal structure of the brain and incompatible with the informational requirements of cognition.

## 2. Encoding stimulus properties

### 2.1. Encoding an experimental parameter

The activity of neurons is often said to encode properties, for example: “*Many cortical neurons encode variables in the external world via bell-shaped tuning curves”* (Seriès et al., 2004). Here the authors refer to a particular type of experiment, where a parameterized stimulus is presented to an animal and the activity of a neuron is recorded. For example the orientation of a small bar is varied and the activity of a neuron in the primary visual cortex is recorded (Fig. 1B). It is found that orientation and neural activity co-vary, and therefore that the neuron’s firing rate encodes the orientation of the bar in the sense of correspondence. What is the scope of such a proposition?

I will discuss a cartoon example from color perception, used by Francis Crick to warn against the “fallacy of the overwise neuron” (Crick, 1979). Cones are broadly tuned to wavelength (Schnapf et al., 1987): in an experiment where light of different wavelengths is flashed, the amplitude of the transduced current varies systematically with wavelength (Fig. 2A). Thus, the current encodes wavelength, in the technical sense of correspondence: one can recover wavelength from the magnitude of the current. Yet animals or humans with a single functional type of cones are color blind. Why are they color blind if their cones encode color information? This is clear in Fig. 2A: if the current also depends on light intensity, then it does not provide unambiguous information about wavelength. In other words, the cone does not in fact encode wavelength in any general setting, even in the narrow sense of correspondence. The same remark applies to any tuning curve experiment.

**Figure 2.**
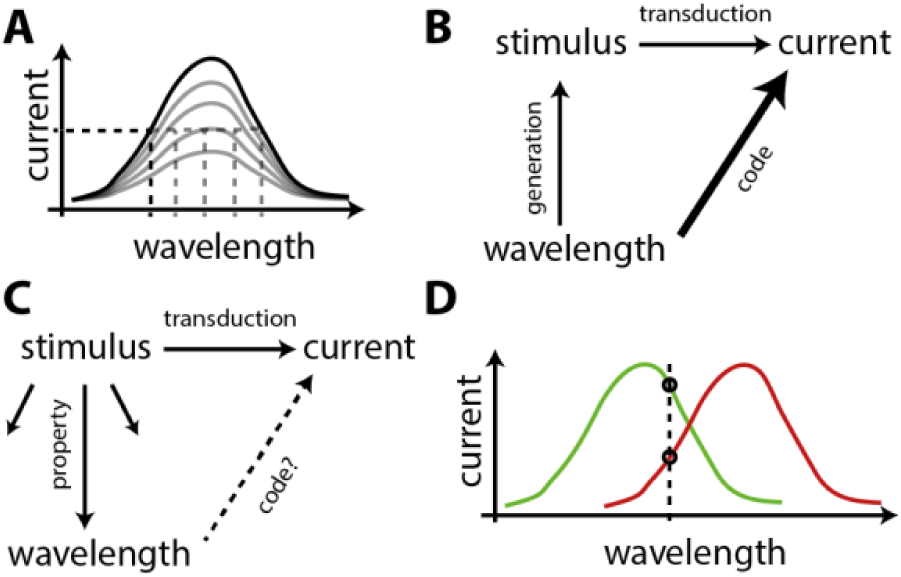
Encoding wavelength of light. A, Response of a cone to flashed light as a function of wavelength (cartoon), at different intensities (grey). If intensity is fixed, wavelength can be inferred from transduced current. Otherwise, current is not informative about wavelength. B, In a tuning curve experiment, the coding relation is implied by the experimental design: wavelength is mapped to stimulus, which is transduced into current. C, If wavelength is just one property of a larger set of stimuli, there might be no coding relation. D, The relative response of cones with different tunings may provide intensity-invariant information about wavelength.

Formally, the logical problem can be analyzed as follows. The tuning curve experiment shows a correspondence between stimulus parameter and current. This correspondence is composed of two parts (Fig. 2B): a mapping from wavelength to stimulus, which is experiment-specific, and the transduction of stimulus into current. Thus, the experimental design ensures that there exists a mapping from wavelength to current. In other words, the proposition that the neuron encodes the experimental parameter is mainly a property of the experimental design rather than of the neuron (which only needs to be sensitive to the parameter). However, the situation is completely different in the real world (Fig. 2C). In general, there might be a variety of stimuli, one of their properties being wavelength. Thus, there is a mapping from stimulus to wavelength and a mapping from stimulus to current, and it is not obvious at all that there is a mapping from wavelength to current, because current depends also on other properties. In this context, the proposition that the neuron encodes wavelength is a much stronger claim, but it is not at all entailed by the tuning curve experiment^1^. This confusion underlies influential neural coding theories of perception, for example Bayesian theories (Pouget et al., 2003; Jazayeri and Movshon, 2006), where a neuron’s firing rate is assumed to be a function of the stimulus parameter, rather than a context-dependent correlate (see section 2.3.2).

Thus, the correct interpretation of the tuning curve experiment is that the neuron is sensitive to the stimulus parameter, while to encode a property of stimuli (a “variable in the external world”) is a somewhat orthogonal proposition: it means that the observable is not sensitive to other properties. For example, a color scientist would point out that wavelength is indeed not encoded by single cones, but by the relative activity of cones with different tunings (Fig. 2D), because that quantity does not depend on light intensity.

Thus, referring to tuning curve experiments in terms of coding promotes a semantic drift, from the modest claim that a neuron is sensitive to some experimental manipulation, to a much stronger claim about the intrinsic representational content of the neuron’s activity. We will now see that this semantic drift indeed operates in current theories of brain function.

### 2.2. The overwise neuron and its ideal observer

To understand how the neural coding metaphor unfolds, I will discuss one particular example in detail (but another one could have been chosen). In mammals, the major cue for sound localization in the horizontal plane is the difference in arrival times of the sound wave at the two ears (interaural time difference or ITD) (Fig. 3A). Neurons in the medial superior olive (MSO) (in the auditory brainstem) are sensitive to this cue (Joris et al., 1998): when a sound is played through earphones and the ITD is varied, the firing rate of those neurons changes (Fig. 3B). These neurons project to neurons in the inferior colliculus (IC), which also have diverse ITD tuning properties. Electrical stimulation in the cat’s IC triggers an orienting response towards a particular contralateral direction, with stronger stimulations resulting in responses with a larger part of the body (one pinna, both pinnae, and eyes), by a pathway involving the superior colliculus (Syka and Straschill, 1970). Unilateral lesions in the MSO or IC result in sound localization deficits in the contralateral field (Jenkins and Masterton, 1982). Thus, neurons in the IC have a critical role in localizing sounds in the contralateral field.

**Figure 3.**
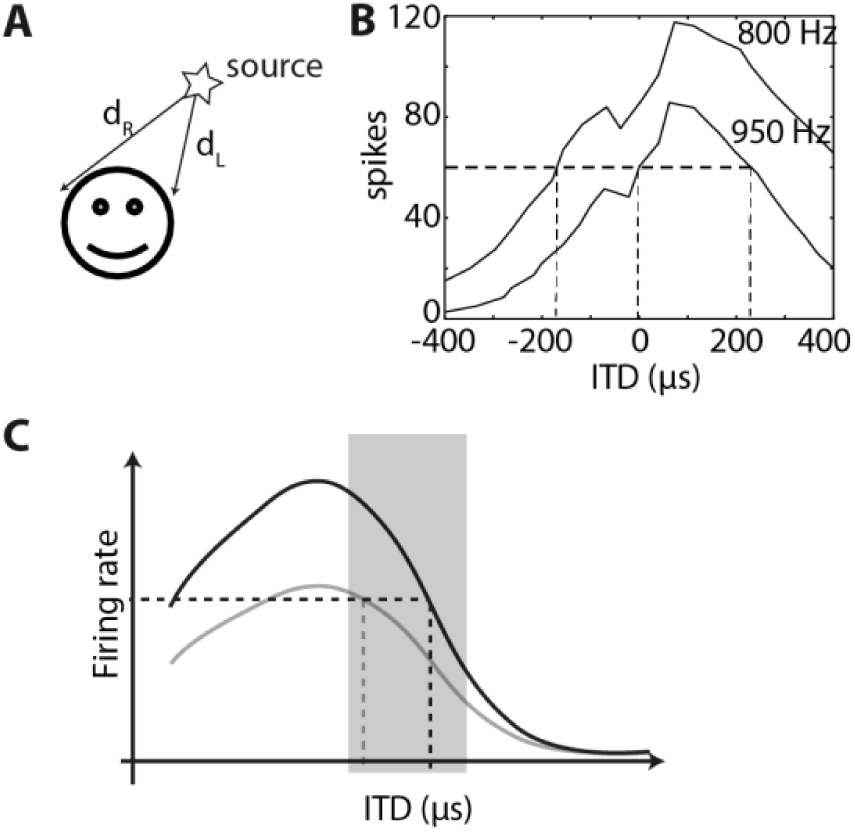
Encoding sound location. A, A major cue for sound localization is the interaural time difference or ITD, d_R_ - d_L_. B, Number of spikes in response to two binaural tones (950 Hz and 800 Hz) as a function of ITD, for the same neuron in the medial superior olive of a cat (digitized from (Yin and Chan, 1990), Fig. 10). It is possible to infer ITD from spike count if the experimental configuration (presented tone) is known, not if the sound is a priori unknown. C, If the organism lived in a world with a single sound played at different ITDs, then the best way to encode ITD would be with a neuron tuned to an ITD outside the range of natural sounds (shaded), so that the selectivity curve is steep inside that range. However, the response of a single neuron is fundamentally ambiguous when sounds are diverse, irrespective of the steepness of the curve (selectivity curve for another sound shown in grey).

How does the activity of these neurons contribute to sound localization behavior? One way is consider the entire pathway and try to build a model of how neuron responses in various structures combine to produce an orientation reflex to a localized sound, and compare with the diverse experimental observations mentioned above. Another way is to ask how neurons encode sound location (McAlpine et al., 2001). It has been claimed for example that *“there is sufficient information in the firing rates of individual neurons to produce ITD just-noticeable-differences that are comparable with those of humans psychophysically”* (Skottun, 1998; Shackleton et al., 2003). What does this mean, and how significant is this fact?

The neuron of Fig. 3B encodes ITD, in the technical sense that one can estimate the ITD with some accuracy from the observation of the number of spikes, by inverting the tuning curve (i.e., decoding the neuron’s response). It turns out that this accuracy is similar to the accuracy of sound localization by the animal. But the neuron’s response is also sensitive to various aspects of sound (e.g. frequency, intensity), so our decoder would give totally inaccurate results in any other context. Thus, the performance of this decoder is unrelated to our general ability to localize sounds. Yet, although the problem of ambiguities was acknowledged, it was concluded that *“it might not be necessary to pool the outputs from many neurons to account for the high accuracy with which human observers can localize sounds’* (Skottun, 1998). This conclusion is unwarranted because tuning curves address the exactly orthogonal problem (sensitivity to ITD vs. insensitivity to other dimensions).

This incorrect conclusion is about localization, but what about discrimination? The first quote compared the tuning curve to discrimination performance, i.e., psychophysical measurements of the ability to discriminate between two sounds that differ only by their ITD. This is a more restricted situation, but how can the responses of a single neuron be compared to the behavior of an organism, without making any reference to the mechanisms that might link this neuron’s activity to behavior? (e.g. the pathway mentioned above). More generally, how can a neural code be about behavior, when it is technically only about stimulus-response properties? This requires what Teller (1984) called a “linking proposition”, an implicit postulate that directly relates neural activity to behavior. The linking proposition here, as in many neural coding studies, is that the brain implements an “ideal observer” (Macmillan and Creelman, 2005). This is the representational sense of the metaphor, namely the idea that neural responses are messages for a reader. The empirical question, then, is how plausible is this linking proposition?

Let us spell it out. The ideal observer reads the activity of the neuron. When the first stimulus is presented, it stores the number of spikes produced by the neuron in a window of a given duration (chosen by the experimenter) after the stimulus. It ignores all spikes produced before and after that window, until the second stimulus is presented, upon which it stores again the number of spikes produced by the neuron. Then it retrieves the two stored numbers, compares them (and not others, e.g. the activity of other neurons), and decides to push one of two buttons. It is not so obvious how this ideal observer can be mapped to the pathway described above, for example how the number of spikes of an arbitrary neuron in the brainstem, produced during several predefined time windows, can be stored in working memory for later comparison.

The ideal observer is ideal in the sense that it makes the best use of all available information. This includes the neural activity itself, but most importantly all the information that is available to the experimenter: when exactly the activity corresponds to the stimulus, what stimulus has been presented, the knowledge that the exact same sounds are played twice, which parts of the activity should be stored. On the other hand, the observer is not ideal in the sense that it uses nothing *more* than the information available to the experimenter. For example, if it also used the information available in other neurons (not recorded), then discrimination performance would be much better than psychophysical measurements. In other words, the ideal observer is not the best thing that the brain can do; it is the best thing that the experimenter can do.

Thus, by implying that the brain reads the neural code, we manage to make claims about perception and behavior while totally ignoring the mechanisms by which behavior is produced, as well as the constraints that the organism must face in ecological situations (e.g. not knowing the sound presentation protocol in advance). These claims rely on implicit linking propositions based on abstract constructs, where neural activity is likened to a processor register that the brain manages to store, retrieve and manipulate, wherever it is in the brain and whenever it occurs. It would seem that empirical evidence or argumentation should be required to support such questionable hypotheses, since all conclusions are based on them. Why is it that no such justification is ever provided when “ideal observers” are introduced? The reason, it seems, is the semantic drift from the technical sense of code to the representational sense of code, which is logically flawed. The same flaw appears to underlie leading theories of neural population coding.

### 2.3. Populations of overwise neurons

#### 2.3.1. Slope coding

What is the optimal way to encode ITD in the activity of neural populations? If all confounding dimensions (level, frequency, etc) are neglected, then the best way to encode ITD is to have a steep monotonous relation between ITD and firing rate, that is, to maximize neural sensitivity to ITD (Fig. 3C). Thus, the neuron’s preferred ITD should lie outside the range of natural sounds (around ±800 μs for humans (Benichoux et al., 2016)) while the steepest slope of the selectivity curve should be inside. This is the concept of “slope coding”. Thus, it has been argued that the optimal way to encode ITD is with two homogeneous populations of neurons with symmetrical tuning curves, peaking at ITDs that are not normally experienced (Harper and McAlpine, 2004). Unfortunately, this conclusion is entirely based on the fallacy of the overwise neuron. If confounding dimensions are not neglected, then the opposite conclusion follows: as in the case of cones, heterogeneity of ITD tunings is crucial to resolve the ambiguities due to non-spatial dimensions of sounds (Brette, 2010; Goodman et al., 2013).

Based on the slope coding idea, a leading theory of sound localization (Grothe et al., 2010) proposes that sound location is encoded in the relative average activity of the two populations of neurons. It was initially meant to explain why many neurons are tuned to large ITDs that are not normally experienced^2^ (McAlpine et al., 2001). Although using the relative activity of two populations somewhat reduces the ambiguity due to sound level, the fact that sounds have more than two dimensions again means that this model is unlikely to work in practice unless the auditory world consists of pure tones (Goodman et al., 2013). In any case, what needs to be demonstrated to support this theory is not that tuning curves have a steep slope, but that the relative average activity of the two neural populations is insensitive to other properties than ITD (e.g. the sound at the source).

Thus the application of the coding metaphor to tuning curve experiments leads to a confusion between parameter sensitivity and information about the corresponding property in a broader context. It could be argued that information in a broad context at least requires sensitivity, but this is also technically incorrect^3^ (see e.g. Zylberberg (2018)).

#### 2.3.2. Encoding visual stimuli

In visual neuroscience, theories of neural coding are rather based on heterogeneous tunings. There are several theories of population coding of stimulus properties in the visual cortex (Pouget et al., 2003; Jazayeri and Movshon, 2006). One influential theory, the “Bayesian brain” hypothesis (Knill and Pouget, 2004), postulates that neural activity represents the probability distribution of the stimulus property, which the brain can manipulate to perform statistical inference. A key assumption in this and other coding theories is that the firing rate of neurons is a context-free function of stimulus properties. This assumption appears explicitly in the models, and in the way the brain is proposed to compute with those representations. For example, one variation of this theory proposes that the brain computes the log likelihood of a stimulus property by summing the activity of neurons weighted by the logarithm of each neuron’s tuning curve (Jazayeri and Movshon, 2006). This operation is described as a *“simple neural readout strategy”*, because it only involves summation and multiplication by fixed weights. As already discussed, the problem is that in reality, tuning curves are defined for a specific experimental condition; they are not context-free. Therefore, either the computation of the log likelihood will be systematically incorrect for all other conditions, or the weights used in the readout must be adapted to correspond to the tuning curve of each condition by an undescribed mechanism, in which case the readout cannot possibly be described as a “simple neural readout”.

To what extent do tuning curves depend on context? As it turns out, to a large extent. It has been known for a long time that properties of sensory neurons adapt to input statistics (Barlow et al., n.d.; Hosoya et al., 2005). In the primary visual cortex, responses to local orientation depend on the surrounding context (Hubel and Wiesel, 1968; Bolz and Gilbert, 1986). Tuning properties of visual cortical neurons (not just the gain) depend on cognitive context, including the task the animal is doing (Gilbert and Li, 2013), locomotion (Pakan et al., 2018) and prior presentation of sounds (Chanauria et al., 2018). Thus, current evidence indicates that the activity of neurons is sensitive to stimulus properties (the technical sense of coding) but cannot be considered as context-free symbols that stand for the corresponding properties (the representational sense of coding). Can neural coding theories of perception accommodate for this fact? It would require that in every context, changes in encoding (stimulus-response properties) are exactly mirrored by changes in decoding (computations performed on neural activity, e.g. the “simple neural readout”). No mechanism has been proposed to achieve this (see also next section).

Theories of neural coding have the ambition to explain some aspects of perceptual behavior, namely results of psychophysical experiments. Again, this requires that a link is made between the neural code and behavior. This link involves ideal observers: for each possible task there is an optimal way to decode neural activity into the variable of interest, which uses detailed elements of the experimental design. Critically, this link with behavior is not considered part of the model because it is assumed that it belongs to the reader of the neural codes^4^. Thus, the behavioral predictions of the coding theories critically rely on linking propositions whose validity or plausibility is not addressed. To be clear, the questionable assumption is not so much whether behavior or perception is optimal in some way (Rahnev and Denison, 2018), but whether the activity of a neuron is something that is read and manipulated as if it were a register of a processor, and not just something that the neuron is doing at a particular time (acting on other neurons).

When the brain is engaged in solving a particular visual task, the activity of neurons depends specifically on object properties relevant to that task (Gilbert and Li, 2013). This seems entirely logical if we see neurons as collaborating to solve a task. In contrast, it is surprising if we see the visual cortex as encoding the world, and the rest of the brain as dealing with this representation to guide actions. Thus, thinking in terms of coding seems to obscure rather than clarify understanding.

### 2.4. Can neurons encode variables?

It could be objected that the problem of contextual dependence of tuning curves only calls for a minor amendment to the mainstream neural coding theories, which is to consider that contextual variables are encoded too. This would require more complicated decoding schemes, but not fundamentally different theories.

For example, one could propose that populations of cones jointly encode wavelength and intensity, and both can be decoded from the joint activity of cones. But to decode cone activity into wavelength, one must know that a monochromatic light is being presented. In natural experience, light is not monochromatic, it has a continuous spectrum, and the transduced current depends on the convolution of the spectrum of incident light with the absorption spectrum of the photoreceptor. In those cases, cones cannot possibly encode wavelength, even jointly, because there is no such thing as the wavelength of a patch of visual scene. Thus, the activity of cones is not sufficient to infer wavelength. A critical element of context that also needs to be encoded is the fact that a monochromatic light is being presented.

Similarly, a cat’s neuron may encode the orientation of a bar only in conjunction with the information that a bar is being presented. That information does not take the form of a variable, but perhaps of a model of the experiment. But models are not variables, rather, they define variables. Thus, a perceptual scene cannot be represented by a set of variables, because this leaves out what defines variables. This missing aspect corresponds to object formation and scene analysis, two fundamental aspects of perception that are not addressed by coding: there is no object property to be encoded if there is no object.

Consider the Bayesian brain hypothesis: *“the brain represents information probabilistically, by coding and computing with probability density functions”* (Knill and Pouget, 2004). This presupposes that there is a set of predefined variables to which probability is attributed – examples of variables are the position of an object, the orientation of a visual grating. If neurons encode variables, then what encodes the definition of those variables, and what do neurons encode in situations where those variables are not defined? We can imagine that such theories might apply to the representation of eye position, for example, because the eye is always there and its position is always defined. This is not the case of objects of perception in general.

Similarly, influential models of working memory propose that memory items are stored and encoded in the persistent activity of neurons tuned to the underlying stimulus property, for example the spatial position of an object (Constantinidis and Klingberg, 2016). This provides a way to store graded properties, like the position of a visual target or the pitch of a musical note. But suppose there is a neural network in my brain that is storing the number 100. What have I memorized? Clearly not the same piece of information if this number is the area of my apartment in square meters or the height of my son in centimeters. To store the information, one needs not only the number but also what it refers to. Can the persistent activity of tuned neurons store that information? It can if there is a network of neurons tuned to the area of my apartment and another tuned to the height of my son.

Perception and memory cannot just be about encoding stimulus properties because this leaves out the very definition of those properties and of the objects they attach to. But could it be that neurons encode more abstract “internal variables” that somehow describe the external world? Such is the claim of predictive coding (Rao and Ballard, 1999) and related propositions such as the free energy principle (Friston, 2009; Clark, 2013). In these theories, neural coding is described as a statistical inference process, where neurons encode the inferred value of internal variables of a generative model of the inputs, e.g. the retinal image. Technically, this essentially means that the code is a parametric description of the image (like a Fourier transform for example). Described at this technical level, the theory seems to have little to say about perception or behavior. But the intended scope extends as these internal variables are described as the “causes” of the sensory input, and the process of encoding is referred to as “inferring the hidden causes”. The sensory input is caused by things in the world, so an internal variable can only be considered a cause if it is assumed to encode properties of objects in the world just like in the Bayesian brain hypothesis. Again, this is incoherent because no perceptual scene can be fully specified by the properties of its objects; one needs first to define objects and their properties, and these definitions are not conveyed by the variables. “Cause” must then be understood in the strict technical sense of variable of a statistical function, which has little to do with the usual sense of “cause”. Thus, this use of the term “cause” appears to be another case of a metaphoric extension of the technical vocabulary. Quoting Chomsky (1959): *“This creates the illusion of a rigorous scientific theory with a very broad scope, although in fact the terms […] [have] at most a vague similarity of meaning”*. We will come back to predictive coding theory in the next part.

Neurons encode stimulus properties according to the technical sense of the metaphor. To acquire a broad scope, the metaphor drifts into the representational sense, according to which neurons convey information about the said properties to the rest of the brain. But neural activity can only be interpreted as properties once the interpretative framework is provided. Critically, this framework is not contained in the coding variables. In what sense do neural codes constitute information for the brain, if their meaning lies outside the encoded messages and varies depending on situations? Where do ideal observers obtain the information necessary to decode the messages? In the next section, I will argue that the coding metaphor conveys a very particular notion of information, which is information by reference, and that this is not the kind of information relevant to perception and behavior.

## 3. Do neural codes constitute information about the world?

### 3.1. Codes as information by reference

The coding metaphor assumes that neural codes represent information about the world, which the brain uses to produce adapted behavior. This sense is implied by the use of ideal observers in the neural coding literature, and more generally by the presumption that the brain “decodes” neural responses or “extracts information” from them. In what sense is the neural code “information” about objective properties of the world? According to the technical sense of coding, it is information in the sense that these properties can be inferred from neural activity. Methodologically, this inference is done by the experimenter, who confronts these properties with measurements of neural activity. But by using the terms “neural code”, and by comparing the output of ideal observers to psychophysical measurements, we imply that the brain must also do this inference.

This raises the issue of “the view from inside the box” (Clark, 2013): how is it possible for the nervous system to infer external properties from neural activity, if all that it ever gets to observe is that activity? In fact, what does it even mean that a neural network infers external properties (e.g. the direction of a sound source), given that those properties do not belong to the domain of neural activity? This is related to the *symbol grounding problem* (Harnad, 1990): how do spikes, the symbols of the neural code, make sense for the organism?

A fundamental issue with the coding metaphor, as it applies to the brain, is that it conveys a very particular notion of information, information by reference: the meaning of the encoded message is that of the original message to which it refers. Shannon made this very clear when he defined his mathematical notion of information (Shannon, 1948):

> *“The fundamental problem of communication is that of reproducing at one point either exactly or approximately a message selected at another point. Frequently the messages have meaning; that is they refer to or are correlated according to some system with certain physical or conceptual entities. These semantic aspects of communication are irrelevant to the engineering problem.”*

But the semantic aspects are precisely what is relevant to the biological problem: how does the brain know what the codes refer to?

One possibility is that the meaning of neural codes is implicit in the structure of the brain that reads them: the brain understands neural codes because it has evolved to do so. There are at least two objections that makes this proposition implausible. First, there is considerable plasticity, including developmental plasticity, both in the nervous system and in the body, which makes the idea of a fixed code implausible. An impressive example is the case of a patient born with a single brain hemisphere, who has normal vision in both hemifields, with a complete reorganization of brain structure (Muckli et al., 2009). This plasticity implies that the “reader” of neural codes must learn their meaning, at least to some extent. Second, one might imagine that the meaning of a neural code for eye position might be fixed by evolution, for example that there are fixed motor circuits that control the eye based on that fixed neural code. But how could this be true of a neural code for the memory item “my apartment is 100 meters square”?

To see that Shannon information cannot be the relevant notion of information to understand perception, consider the following experiment of thought, which I shall call the paradox of efficient coding. Suppose that all information about the world (including efferent copies) is encoded by a set of neurons. From the heuristic that biological organisms tend to be efficient, we now postulate that neurons transform their inputs in such a way as to transmit the maximum amount of information about the world, in the sense of Shannon; this is the efficient coding hypothesis (Barlow, 1961; Olshausen and Field, 2004). This means that all redundancy is removed from the original signals. If this is done perfectly, then encoded messages are undistinguishable from random. Therefore the perfectly efficient code cannot be understood by its reader.

It is indeed paradoxical that when we maximize the amount of information carried by code, we find something that provides no information at all to the reader. This is because the notion of information implied by the phrase “neurons encode information” is information by reference to the inputs, a kind of information that is only accessible to an external observer. This is not the right way to address the informational problems faced by the brain. Again, the coding metaphor appears to promote a semantic drift, from the technical sense of information as defined by Shannon, to a broader sense of information that might be useful for an organism. The neural coding metaphor is so prevalent in the neuroscience literature that the notion of information it carries seems to be the only possible one: *“the abstract definition of information is well motivated, unique, and most certainly relevant to the brain”* (Simoncelli, 2003). Next, I discuss alternative notions of information that are more relevant to the brain.

### 3.2. Information as subjective laws or internal models

How can there be any information about the world without direct access to the world? John Eccles, a prominent neurophysiologist, expressed the problem in the following terms (Eccles, 1965):

> *“In response to sensory stimulation, I experience a private perceptual world which must be regarded, neurophysiologically, as an interpretation of specific events in my brain. Hence I am confronted by the problem: how can these diverse cerebral patterns of activity give me valid pictures of the external world?”*

To him, the logical solution was a form of dualism, much like Cartesian dualism, except he did not believe that the interaction between mind and brain occurred at a single place (Descartes’ pineal gland). Dualism is a natural solution if neural activity is thought to encode information by reference to the external world, because the external world belongs to a different domain.

A number of philosophers and psychologists have proposed alternative solutions. O’Regan and Noë (2001) proposed the analogy of the “villainous monster”. Imagine you are exploring the sea with an underwater vessel. But a villainous monster mixes all the cables and so all the sensors and actuators are now related to the external world in a new way. How can you know anything about the world? The only way is to analyze the structure of sensor data and their relationships with actions that you can perform. If dualism is rejected, then this is the kind of information that is available to the nervous system. A salient feature of this notion of information is that, in contrast with Shannon’s information, it is defined as relations or logical propositions: if I do action A, then sensory property B happens; if sensory property A happens, then another property B will happen next; if I do action A in sensory context B, then C happens.

James Gibson previously developed a related psychological theory (Gibson, 1986). While criticizing the information-processing view of perception, he argued that there is information about the world present in the *invariant structure* of sensory signals: “*A great many properties of the [optical] array are lawfully or regularly variant with change of observation point, and this means that in each case a property defined by the law is invariant”*. Clearly, he did not mean information in the sense of communication theory, but rather in the sense of scientific knowledge. A set of observations and experiments provides information about the world, in the form of laws that relate observables (sensory signals) between them and with possible actions. This form of information is intrinsic; I proposed to call this set of laws the subjective physics of the world (Brette, 2016). A related view, formalized by theoretical biologist Robert Rosen (Rosen, 1985), is that biological organisms build an internal model of the world, in which the variables are sensory signals. This view addresses the symbol grounding problem by mapping sensory signals to elements of an internal model. The signals make sense in reference to that model; they are not mapped to externally defined properties.

Crucially, relations between observables are precisely what neural coding theory considers as redundancy, which ideally should be eliminated. In contrast, in the alternative view discussed here, relations constitute information. This point was made by Thompson (1968). *“It is our subjective habit to organize the individual elements of our experience, to cross-correlate these elements to others distant in space and time, and it is only after this process of imposing organization that we feel informed”*. The number 100 does not really constitute information; it is only once I have inserted it into my internal model of the world by saying that it is the area of my apartment in square meters that it becomes information.

### 3.3. Subjective physics of the Martian iguana

To make this point more concrete, I will discuss an example adapted from Brette (2016). Consider a fictional organism with two ears – let us call it a Martian iguana in reference to Dennett (1978) (Fig. 4A). The iguana is fixed on the ground, and there is another organism – let us call it a frog – which produces sounds. The frog is usually still and produces some random sounds repeatedly, but occasionally it jumps to a new position. The question is: what kind of information can the iguana have access to, based on the acoustical signals at the two ears?

When a source produces a sound, two sound waves SL and SR arrive at the two ears, and these two sound waves have a particular property: they are delayed versions of each other (S_L_(t) = S_R_(t-Δ)) (Fig. 4B). In Gibsonian terminology, there is “invariant structure” in the sensory flow, which is to say that the signals obey a particular law. Thus, the sensory world of the iguana is made of random pairs of signals which follow particular laws that the iguana can identify. This identification is what Gibson called the “pick-up of information”. Evidently, “information” is not meant in the sense of Shannon but in the sense of laws or models of the sensory input. Note that the model in question is not a generative model as in predictive coding, but relations between observables, like the models of physics.

A first interesting aspect of this alternative notion of information is that the topology of the world projects to the topology of sensory laws. By this, I mean that two different sounds produced by the frog at the same position will produce pairs of signals (S_L_, S_R_) that share the same property (the sensory law). This can be assessed without knowing what this property corresponds to in the world (i.e., the frog’s position).

Thus, the iguana can observe sensory laws that have some particular properties, but do these laws convey any information about where the frog is? For an external observer, they certainly do, since the delay Δ is lawfully related to the frog’s position. For the iguana, however, they do not because that lawful relation cannot be inferred from just observing the acoustical signals. Thus, this organism cannot have any sense of space, even though neural coding theories would pretend that it does, based on the correspondence between frog position and the activity of the iguana’s auditory neurons.

Let us now consider in addition that the iguana can turn its head (Fig. 4C). It can then observe a lawful relation between a proprioceptive signal (related to the head’s position) and the observed delay Δ, which holds for some time (until the frog jumps to another position). Now when the iguana observes sounds with a particular delay, it can infer that if it were to move its head, then the delay would change in a particular predictable way. For the iguana, the relation between acoustical delay and proprioception *defines* the spatial position of the frog. We note that the perceptual inference involved here does not refer to a property in the external world (frog position), but to manipulations of an internal sensorimotor model.

Thus, the kind of information available to an organism is not Shannon information (correspondence to external properties of the world), but internal sensorimotor models. The interest of such models for the animal is that they can be manipulated so as to predict the effect of hypothetical actions.

**Figure 4.**
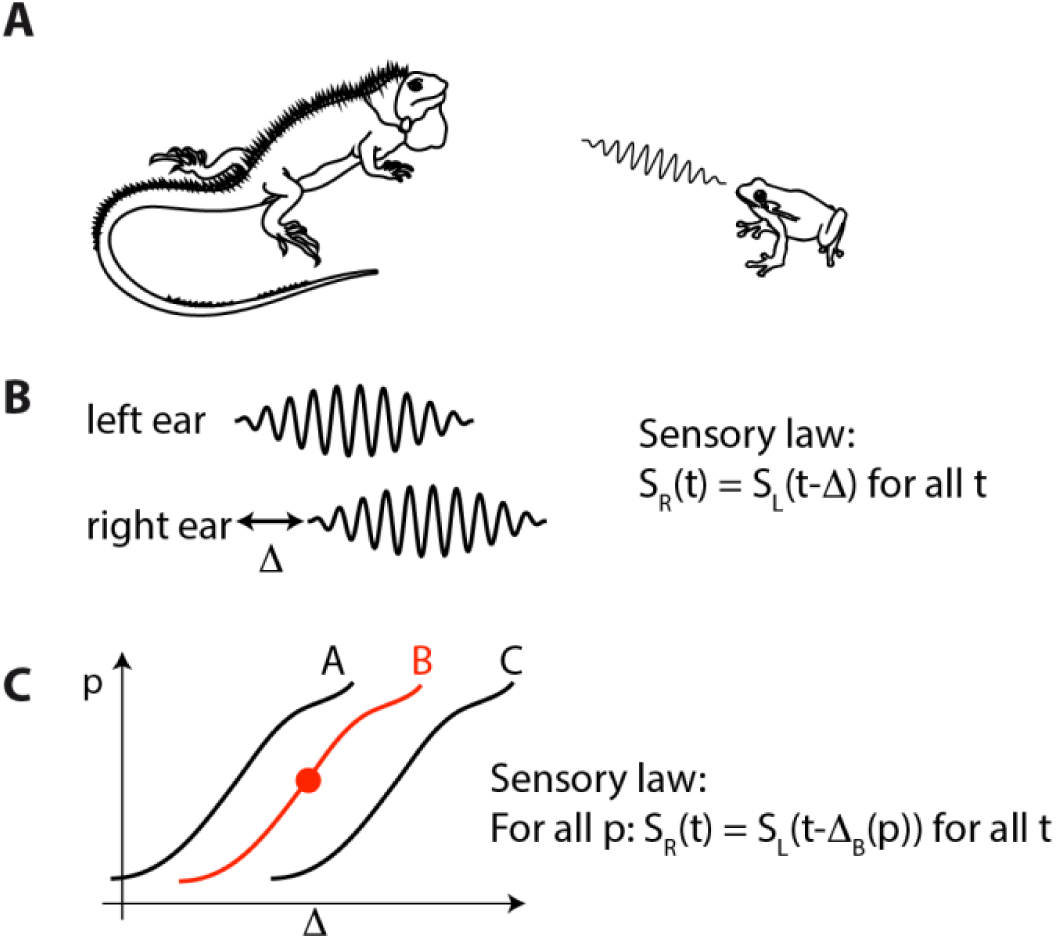
Subjective physics of a fictional iguana. A, The (blind) iguana listens to sounds produced by the frog, which occasionally jumps to a new position. B, When there is a sound, the iguana can notice that the acoustical signals at its two ears follow a particular law: S_R_(t) = S_L_(t-Δ) for all t. C, If the iguana can move its head, it can also notice that the delay Δ changes in a lawful way with the proprioceptive signal p. This relation defines the frog’s position for the iguana. When a sound is heard, the iguana can infer the frog’s position, i.e., it can infer how Δ would change if it were to move its head.

### 3.4. Predictive coding and generative models

Predictive coding theory and its derivatives (Rao and Ballard, 1999; Friston, 2010; Clark, 2013) propose that the brain encodes an internal model, which predicts the sensory inputs^5^. This seems to resemble the proposition of the previous section. More precisely, neurons are thought to encode the variables of a hierarchical model of the inputs, in which higher order neurons encode their prediction of the activity of neurons lower in the hierarchy, down to the sensory inputs. This prediction is subtracted from the input of lower-order neurons, so only the prediction error remains. This leads to a compressed representation of the inputs, and in this sense it is a type of efficient coding theory.

This particular kind of model is called a generative model because its maps internal variables to the observables (sensory inputs), in contrast with the models of physics which take the form of relations between observables (e.g. the ideal gas law, PV = nRT). Generative models are not the kind of internal models described in the previous section.

Consider the iguana with a fixed head. A generative model of the sensory inputs would map two internal variables S (sound) and *Δ* (interaural delay) to the two acoustical inputs S_L_ and S_R_, as: *S_L_(t) = S(t), SR(t) = S(t-Δ)*. Neural activity encodes not the model itself but the coding variables *Δ* and S. In particular, neurons encode the entire sound S, even though it carries no information for the iguana (S is, by construction, random). This appears to contradict the claims of predicting coding theory: *“To successfully represent the world in perception […] depends crucially upon cancelling out sensory prediction error”* (Clark, 2013). Indeed the success of a predictive code is evaluated by its ability to represent the input in a pictorial sense (as if it were a painting), but in this example, the numerical value of the signals provides no useful information beyond the relations they obey.

Consider now the case when the iguana can move its head. The internal model discussed in the previous section is: *S_R_(t) = S_L_(t-Δ_x_(p))* for all t, where x is the frog position (Fig. 4C). The usefulness of this model stems from the fact that it can be manipulated, that is, upon hearing a sound, the iguana can infer that, if it were to move its head to a new position *p*, then the relation obeyed by the auditory signals would change in a predictable way. For example, the iguana can move its head so that S_R_ = S_L_ (“the frog is in front”). Thus, the kind of prediction that this model can produce is about relations between signals, and not about the numerical value of the signals.

On the other hand, a generative model would map the coding variables S, *x* and *p* to the sensory inputs *S_L_(t) = S(t)* and *SR(t) = S(t-Δ_x_(p))*. This mapping is referred to as “prediction”, and is instantiated by the feedback from higher-order neurons to lower-order neurons. This is not the same sense as predicting what action would make the two signals S_L_ and S_R_ match. This brings us to a discussion of the term “predictive” in predictive coding. The appeal of predictive processing is that making predictions seems to be a prerequisite to goal-directed behavior, and thus a fundamental aspect of behavior. In fact, several authors have argued that anticipation is not just a property of nervous systems, but even a fundamental property of life (Maturana and Varela, n.d.; Rosen, 1985). For example, the iguana can predict how some properties of its sensory inputs should change if it were to turn its head. Consider this other example in human behavior: when someone is facing a cliff, she tends to slightly lean backwards, because this posture makes it easier to move backwards if necessary (Le Mouel and Brette, 2017). But this is not at all the technical sense of “prediction” in predictive coding, as remarked by Anderson and Chemero (2013). A neuron “predicts” the sensory inputs in the sense that its firing correlates with them; more specifically, a spike produced by a neuron leads to a subtraction of the expected input of a target neuron, which is the input happening now, or possibly if we incorporate conduction delays, what will be happening after a fixed delay. This is not the kind of prediction implied by an anticipatory postural adjustment: if I change my posture in this way, then it will be easier to move backwards in the hypothetical event that my balance is challenged.

In fact, what is useful for the organism is not literally to predict what will happen next, but rather what *might* happen next, conditionally on the actions I can do, so that I can select the appropriate action. But this requires to manipulate the model. For example, to select an action requires instantiating the internal model with several possible values of action, then to calculate the expected sensory variables. But this contradicts the proposition that neurons encode the “causes” of current sensory signals: to manipulate the model, encoding neurons would then have to be somehow disconnected from the sensory stream.

Technical work on predictive coding has focused exclusively on the technical senses of prediction and coding (correspondence), and thus there is no empirical evidence that such codes might allow the organism to form predictions in a broader sense, nor is there any indication of how a theory based on neural coding might in principle explain anticipatory behavior.

### 3.5. Can neural codes represent structure?

Thus, the kind of representation of the world useful for adapted behavior is a structured internal model. Can neural codes possibly represent that structure? Memories and percepts are thought to be encoded by cell assemblies. In its basic and most popular form, a cell assembly is simply a specific subset of all neurons. When neurons of a cell assembly activate, the corresponding percept is formed (possibly indirectly by the activation of target neurons). This is the basic assumption of associative neural models of memory (Tonegawa et al., 2015): retrieving a memory consists in triggering activity in part of the memory-specific cell assembly (or “engram cells”), which then leads to the activation of all neurons in the assembly.

One problem with cell assemblies, in this simple form, is that they are unstructured, and therefore they cannot represent structured internal models. The cell assembly model is analog to the “bag of words” model in text retrieval, where a text is represented by its set of words and all syntax is discarded. In essence, a cell assembly is a “bag of neurons”. This causes a problem to represent not just the lawful structure of the world, but also the structure of any given perceptual scene. Consider for example the simple visual scene depicted in Fig. 5. There is Paul, a person I know, wearing a new shirt, driving a car (Fig. 5). What is important here is that a scene is not just a “bag of objects”: objects have relationships with each other, and there are many possible different relationships. For example there is a car and there is Paul, and Paul is in a specific relationship with the car, both a physical relationship (a particular posture within the car) and a functional relationship (driving it). Some of my behavior depends on identifying these relations, since for example I can talk about them, and so if behavior relies on neural codes, then those codes should represent relations, not just the pixels of the image.

But cell assemblies cannot represent these relations. Suppose there is a cell assembly that encodes “Paul” and another one that encodes “car”. To encode the driving relation between Paul and the car, one would need a cell assembly that encodes “driving”, but that assembly should also somehow refer to the two assemblies representing Paul and the car, and this is something that cannot be done with an unstructured bag of neurons (mathematically, one would need a labeled graph and not just a subset of nodes).

**Figure 5.**
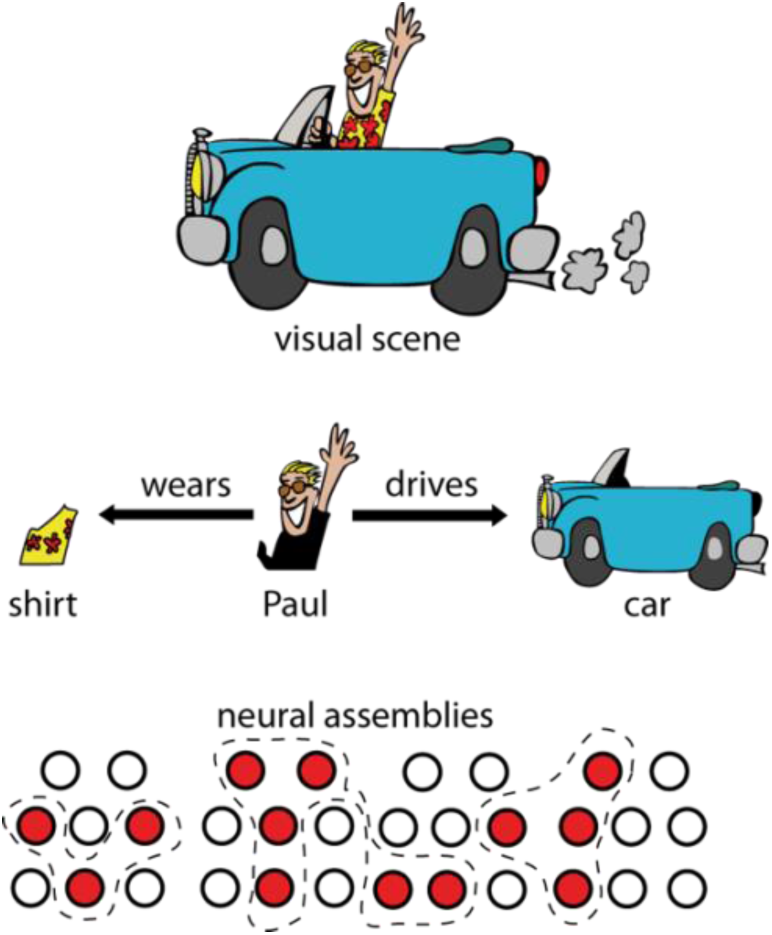
Perceptual scenes are highly structured. For example, there is Paul (person I know), driving a car, and wearing a new shirt. Representing this scene by the firing of neural assemblies raises two issues: 1) it may be difficult to split active neurons into the correct assemblies (superposition catastrophe), and more importantly 2) the structure of the scene (relations shown by arrows) cannot be represented in this way.

This is related to the “binding problem”, although broader. If it is true that any given object is represented by the firing of a given assembly of neurons, then several objects should be represented by the firing of a bigger assembly of neurons, the union of all assemblies, one for each object. Several authors have noted that this may lead to the “superposition catastrophe” (von der Malsburg, 1999), i.e., there may be different sets of objects whose representations are fused into the same big assembly. One proposition is that the binding problem could be solved using retinotopic position as an object label, i.e., neurons do not encode features but the conjunction of feature and retinotopic position (Kawato, 1997). However, this objection does not address the broader point, which is that cell assemblies encode objects or features to be related, but not the relations between them. In fact, it is known that current connectionist models, which are designed to optimally implement the idea that features are represented by the activity of one or several cells, cannot be trained to detect very simple relations between shapes in an image (Ricci et al., 2018).

The binding problem has led several authors to postulate that synchrony is used to bind the features of an object represented by neural firing^6^ (Singer, 1999; von der Malsburg, 1999). This avoids the superposition catastrophe because at a given time, only one object is represented by neural firing. Synchrony is indeed a relation between neurons (mathematically, an equivalence relation). There are a few other examples in the neuroscience literature where synchrony is used to represent relations, although they are not usually cast in this way. One is the Jeffress model of ITD coding (Jeffress, 1948) (Figure 6A). In that model, neurons receive inputs from monaural neurons on the two sides, with different conduction delays. When input spikes arrive simultaneously, the neuron spikes. Thus, the neuron spikes when the two acoustical signals at the two ears are such that S_L_(t) = S_R_(t-d), where d is the conduction delay mismatch between the two ears. Physically, this corresponds to a sound source placed at a position such that it produces an ITD equal to d. In this model, the neuron’s firing indicates whether signals satisfy a particular sensory law.

**Figure 6.**
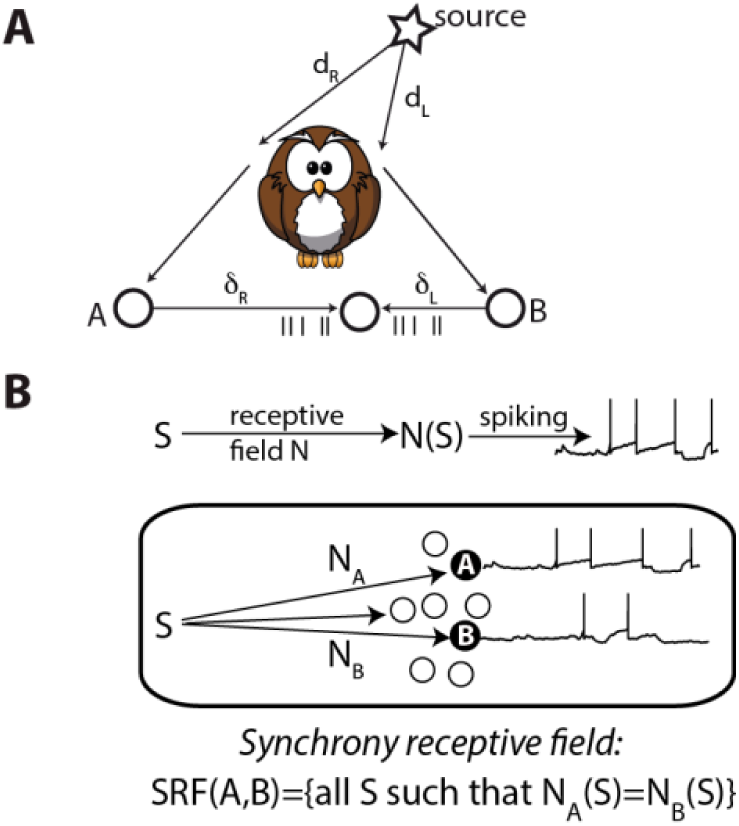
Neural representation of structure (adapted from Brette (2012)). A, The Jeffress model of sound localization. The sound arrives at the two ears with delays d_L_ and d_R_. It is then transduced into spike trains that arrive at a binaural neuron with delays δ_L_ and δ_R_. Synchrony occurs when d_R_ - d_L_ = δ_L_ - δ_R_, making the neuron fire. B, The synchrony receptive field. The response of a neuron to a stimulus is described as filtering of the sensory signal S through the receptive field N, followed by spiking. The synchrony receptive field of two neurons A and B with different receptive fields N_A_ and N_B_ is defined as the set of stimuli that elicit synchronous responses in these neurons.

This interpretation of the model has been generalized with the concept of “synchrony receptive field” (Brette, 2012), which is the set of stimuli that elicit synchronous responses in a given group of neurons (Fig. 5B). One considers two neurons A and B which convert their time-varying inputs into precisely timed spike trains, where their inputs are seen as transformed versions N_A_(S) and N_B_(S) of the stimulus S (N_A_ and N_B_ are fixed and correspond to the receptive fields of the neurons). Synchrony between A and B then reflects (“encodes”) the sensory law N_A_(S) = N_B_(S). This framework has been applied to pitch perception (Laudanski et al., 2014) and to sound localization in realistic environments (Goodman and Brette, 2010; Benichoux et al., 2015).

However, although synchrony can represent relations, neither binding by synchrony nor synchrony receptive fields solves the general problem (even theoretically), because only one type of relation can be represented by synchrony, and a symmetrical one: does Paul drive the car, or does the car run over Paul? The fact that sentences can represent relations motivates the idea that the temporal structure of neural activity (e.g. the sequence of activated neurons, much like a sequence of words) could perhaps provide the adequate basis for structured neural representations (Buzsáki, 2010). But this possibility remains speculative, and in particular it remains to be demonstrated that such hypothetical structures have the quality of representations that the brain can manipulate.

## 4. The causal structure of the coding metaphor

In the previous parts, I have argued that the scope of neural coding theories lies in their use of the representational sense of the metaphor, the idea that neural codes are symbols standing for properties that the brain manipulates, but no evidence has been provided that this sense is valid. Worse, there is empirical evidence and theoretical arguments to the contrary.

Here I focus on a deeper problem with the neural coding metaphor. A striking characteristic of this metaphor is that it is a way to think about the brain independently of its causal structure. When we say for example that neurons encode the location of sounds, we talk about the activity of neurons without making any reference to the result of that activity, or to the system of which the neurons are a component. I now examine the implications of this fact.

### 4.1. The dualistic structure of the coding metaphor

The coding metaphor has a dualistic structure. It structures the function of the brain into two distinct and dual components: the component that encodes the world into the activity of neurons, and the dual component that decodes that activity into the world, or into actions in the world. For example: *“Information that has been coded must at some point be decoded also; One suspects, then, that somewhere within the nervous system there is another interface, or boundary, but not necessarily a geometrical surface, where ‘code’ becomes ‘image.’“* (Somjen, 1972); *“interpretation of the encoded information, typically consisting of its recoding by a higher-order set of neurons or of its “decoding” by an effector”* (Perkel and Bullock, 1968); “A *stimulus activates a population of neurons in various areas of the brain. To guide behavior, the brain must correctly decode this population response and extract the sensory information as reliably as possible.”* (Jazayeri and Movshon, 2006); *“the brain typically makes decisions […] by evaluating the activity of large neuronal populations”* (Quian Quiroga and Panzeri, 2009). “Ideal observers” used in many studies implement this dual decoding brain.

Using the coding metaphor does not necessarily mean believing in dualism of body and mind^7^, but its dualistic structure has important consequences when it comes to understanding function. The two dual components (encoding/decoding) are indistinguishable in behavior, because no behavior involves just one of them. How then is it possible to attribute function to neural codes? How is it possible to draw conclusions about the neural basis of behavior from properties of neural codes, independently of the system in which the neurons are embedded? This is only possible by making an additional assumption, namely that the encoding component has a function by itself (representing the inputs), somehow assigning the status of organ to a part of the nervous system. But there is no indication that the brain can be functionally decoupled in this way: neuroanatomy rather seems to invalidate this hypothesis.

To illustrate this point, I will now discuss a concrete biological example. Paramecium is a unicellular organism that swims in stagnant fresh water using cilia and feeds on bacteria. It uses different kinds of sensory signals, including mechanical signals to avoid obstacles and chemical signals to localize food (Jennings, 1906). To a first approximation, it alternates between straight courses and sudden random changes in direction (Fig. 7A). It turns out that each change in direction is triggered by a spike produced by voltage-gated calcium channels (Fig. 7B) (Eckert, 1972). To find a chemical source, Paramecium uses a simple method: when concentration decreases, the membrane is depolarized by chemical receptors and a spike is produced (with some stochasticity), triggering a change of direction (similar to chemotaxis in E. Coli). This is of course a simplified description of Paramecium physiology and behavior, but for the sake of the argument we shall consider an organism that functions in this simple way.

**Figure 7.**
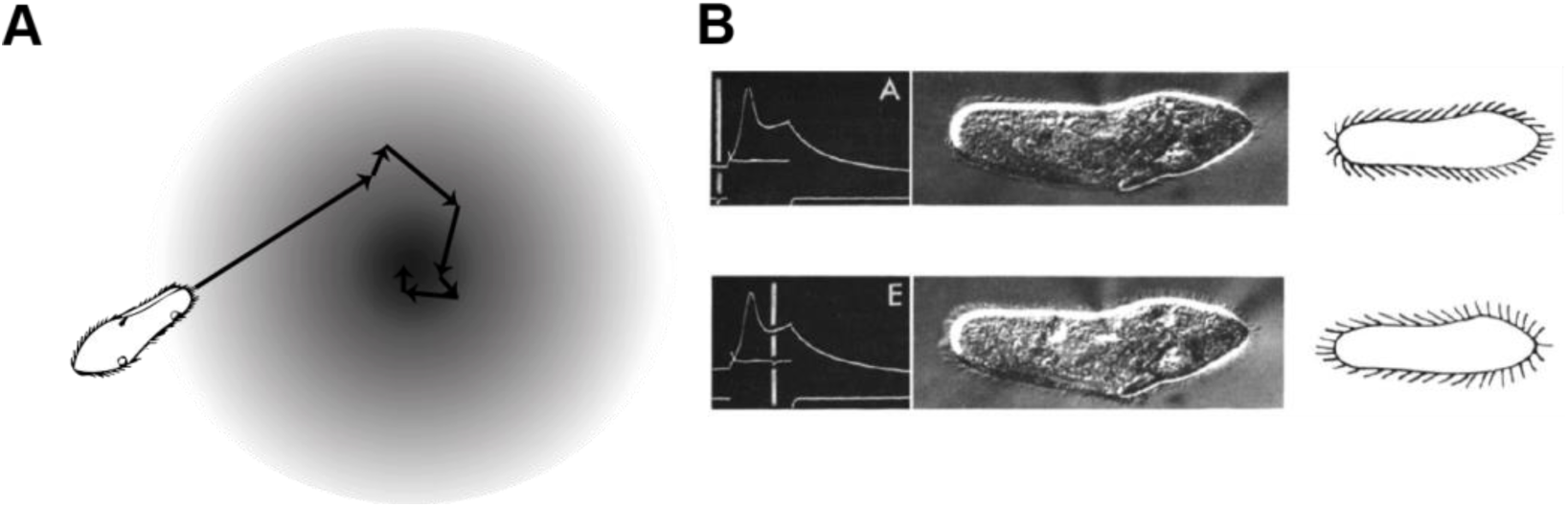
Spatial cognition in Paramecium, a “swimming neuron”. A, Paramecium finds a chemical source by switching to a new random direction when concentration decreases. B, Each direction change is triggered by an action potential, which transiently inverts cilia beating through a calcium pathway (adapted from (Eckert and Naitoh, 1970)).

Thus, Paramecium is a sort of swimming neuron. Spiking activity varies lawfully with sensory signals (concentration), and therefore encodes them in the same sense as a visual cortical neuron encodes visual signals. Just as for sensory neurons of the brain, we may argue that if the organism can navigate efficiently in its environment, then the spikes must contain information about that environment. Thus, it seems that the coding metaphor applies equally well to this swimming neuron as to any typical case in neuroscience.

Let us know think about functional questions. As an organism, Paramecium may have goals, for example to find food. We may hypothesize that it achieves this goal efficiently, for example by finding food as quickly as possible. To this end, sensory signals must be transformed into spikes in a specific way, which depends both on the goal (to move towards or away from a source, to look for food or to sleep, or to look for a mate) and on the effect of spikes on the organism’s actions. Thus, there is a way to organize this system so that it achieves its function appropriately, which determines the transformation of inputs into spikes, i.e., the neural code.

But if we now think of the neural code independently of the organism and environment that host it, we draw different conclusions. If the function of this neuron is to encode its input, then we may hypothesize that it achieves this function efficiently. This prescription determines a neural code that is specified by the statistics of inputs. Here the code depends neither on the goals of the animals nor on the effect of spikes on the organism’s actions. It follows that this efficient code does not match, in general, the neural code that is adapted for the organism’s goal. This mismatch occurs because function can be meaningfully ascribed to the organism as a system, but not necessarily to the components of this system.

This sensorimotor system is arguably much simpler than the brain, nevertheless it demonstrates that the function of neurons cannot be meaningfully framed in terms of coding just because they respond to sensory stimuli. There is no indication that the brain is special in that it can be meaningfully separated into two dual components with independent functionality.

### 4.2. Coding vs. causing

The Paramecium example highlights the fact that the neural coding metaphor is a way to think about the brain that is disconnected from its causal structure. Yet by postulating that neural codes are representations, we imply that these codes have a causal impact on the brain. This is also the case when neural codes are considered simply as transformations of inputs rather than explicit representations, as in Perkel and Bullock (1968): *“The problem of neural coding is defined as that of elucidating the transformations of information in the nervous system, from receptors through internuncials to motor neurons to effectors”*. But does coding implies causing?

Consider for example the BOLD signal, a property of blood used for functional brain imaging because it covaries with neural activity. Thus, the BOLD signal encodes visual signals in the same technical sense that the firing of neurons encodes visual signals. For example, one can “decode” the image from this signal (Naselaris et al., 2009). Yet, visual perception is not caused by the BOLD signal, which is why we do not consider that it is an internal representation used by the brain. Thus, not all coding variables have causal powers.

Consider the firing rate vs. spike timing debate (Kumar et al., 2010; Brette, 2015). This debate is generally formulated in the following way: “does the brain use a firing rate code or a spike timing code?”. As the previous example illustrates, this is a largely irrelevant question because it focuses on correlations between stimuli and observables. We may as well ask: “does the brain use the BOLD code?”. The relevant question is rather whether those observables have a causal role in the activity of the brain, and this involves a different set of arguments and answers (see Brette (2015) for a discussion). To see why, consider a sensorimotor system whose function is well understood in relation to its electrical activity: the heart (Fig. 8). The heart operates like a pump to circulate blood in two phases: the two atria contract, pushing blood into the ventricles (diastole); then the two ventricles contract, pushing blood into the pulmonary arteries (systole). These contractions are triggered by excitable cells in atria and ventricles. For the heart to operate as a pump, cells in the two atria must spike synchronously, but out of phase with cells in the two ventricles. But the heart also responds to sensory stimulation. For example, the heart beats at a faster pace when we run. This means that the excitable cells of the heart encode running speed in their firing rate, in the technical sense. If we now look at coding properties of these cells, we find that: 1) firing rate is sensitive to running speed, 2) cells fire regularly, 3) spike timing is not reproducible between trials, 4) spike timing (absolute or relative) carries no information about the stimulus beyond the rate. Thus, we would conclude that the heart uses a rate code. Yet, the temporal coordination of spikes is critical in this system; in fact, it is life-critical. This paradox arises because the neural coding metaphor totally neglects the causal effect of spikes.

**Figure 8.**
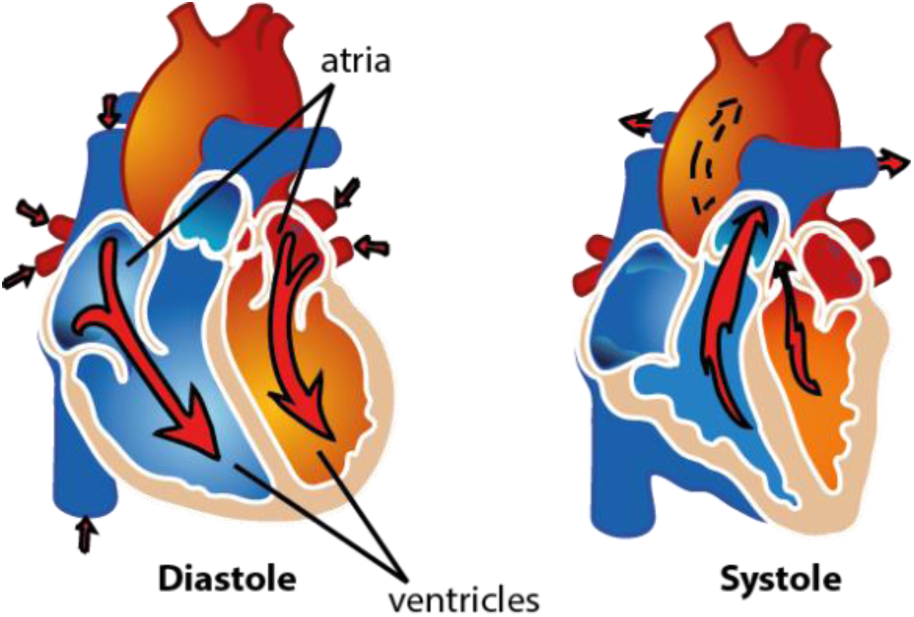
Operation of the heart: atria simultaneously contract, triggered by synchronous firing of excitable cells, then ventricles simultaneously contract, pushing blood into the lungs.

Thus, if we want to describe the operation of the brain in terms of neural coding, the relevant question is whether the causal structure of neural codes is congruent with the causal structure of the brain. If it is not the case, then the neural coding metaphor loses much of its scope.

### 4.3. Causal powers of neural codes

The causal structure of the brain is sketched on Figure 9A. At a coarse description level, the brain is a dynamical system coupled to the environment by circular causality. At a finer description level, the brain is itself made of neurons, which are themselves dynamical systems coupled together. To a first approximation, the coupling is mediated by spikes, which are timed events. In a dynamical system, state variables have a causal role by construction; examples of state variables in this physical system are membrane potential and the state of ionic channels. Spikes have causal effect, but being events they are not state variables; a spike is something that happens, not a property of the system.

**Figure 9.**
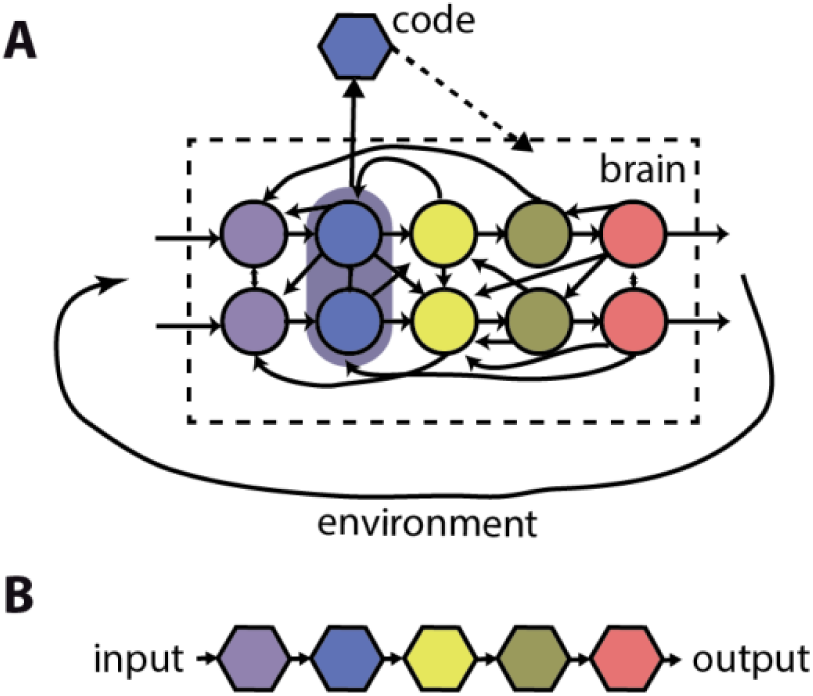
Causal structure of brain and neural codes. A, The brain is a distributed dynamical system made of interacting neurons, which is coupled to the environment by circular causality. A code is a property of neural activity, which is implicitly assumed to have a causal effect on the brain. B, Neural codes are linked together and with the world by linear causality.

Let us now examine the causal structure of neural codes. A neural code is a correspondence between sensory inputs (or an external property) and a coding variable. The coding variable is an aggregate variable based on measurements of spiking activity over some time, space and possibly trials. Aggregate variables can have causal effect in principle, for example order parameters in statistical physics, which are macroscopic variables that determine phase transitions (e.g. ice melting). But of course not all observables are order parameters. What about coding variables? I will consider integrating over trials, space and time in turn.

An example of integrating over trials (and time) is a neuron that responds specifically to pictures of Jennifer Anniston in various poses (Quiroga et al., 2005). But only on average: the coding variable is the median number of spikes across trials between 300 and 1,000 ms after stimulus onset. On a given trial, the neuron might not be firing at all. Unless the subject was not perceiving the actress in those trials, this implies that this neuron cannot encode the percept “Jennifer Anniston” in the sense of causing the percept. Rather, its firing correlates (on average) with the presentation of Jennifer Anniston pictures – which is already a notable fact. Perceptual representations cannot be based on averages; percepts are experienced now, not on average. Neural codes based on averaging over trials do not have causal powers. In the same way, a firing probability (one abstract way to define a neuron’s firing rate) does not have causal powers^8^; only the occurrence of firing does.

An example of integrating over space (and time) is when we propose that the position of a sound source is encoded by the difference in total activity between the two symmetrical inferior colliculi (Grothe et al., 2010). This coding variable indeed varies when source position is changed (Thompson et al., 2006). Does it mean that it has causal powers, i.e., that it determines sound localization behavior? It seems implausible, first as previously discussed because it also varies with other properties of sounds, second because electrical stimulation in the inferior colliculus triggers orienting responses which vary with the place of stimulation, while stronger stimulation results in orienting responses that engage a larger part of the body (one pinna, both pinnae, and eyes, in order of recruitment) (Syka and Straschill, 1970). Thus, there is no guarantee that a coding variable obtained by integrating over neurons has causal powers.

But the key difficulty is time. In an experimental setting, the coding variable is obtained by measuring spiking activity over a certain duration of observation, following the onset of the stimulus. Thus, it is tied to the temporality of the experiment. But to have any causal power, the code must be somehow related to the temporality of the system with which it interacts. The brain does not wait for one second after stimulus onset until it processes the activity of some neurons. The brain has an ongoing spontaneous activity that is similar in many ways to stimulus-driven activity (Deco et al., 2011). It is influenced by sensory signals, which have no definite onset. Finally neurons mutually influence themselves over timescales of a few milliseconds, without waiting for the coding variable to be defined. Neural codes abstract time away, but temporality is critical to the operation of a dynamical system.

The neural coding metaphor leads to a description of the operation of the brain as a sequence of neural codes (Fig. 9B), as in Perkel and Bullock’s (1968) definition of neural coding (*“transformations […] from receptors through internuncials to motor neurons to effectors”*). In this description, the temporality of the physical system has disappeared, and has been replaced by the discrete temporality of an algorithm, which is disconnected from physical time. In other words, this is an algorithmic description. But as van Gelder (1995) pointed out, dynamical systems cannot in general be mapped to algorithmic descriptions^9^. For the coding view to be valid, it must be demonstrated that the dynamical causal structure of Fig. 9A can be fitted into the linear algorithmic structure of Fig. 9B. But this demonstration has never been provided. What has been empirically demonstrated is that the input is in correspondence with increasingly complex codes, with coding variables measured over extended time windows tied to stimulus-response experiments.

**Figure 10.**
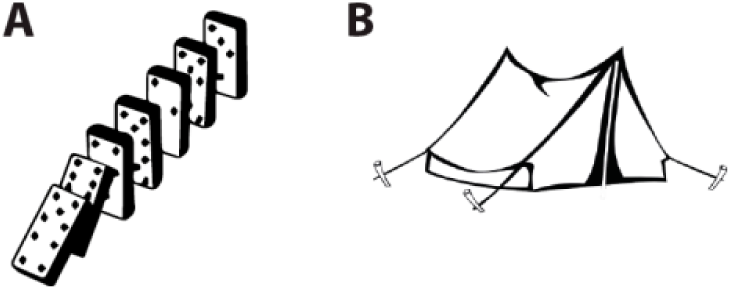
The causal structure of neural coding metaphor is that of dominoes (A), but the causal structure of the brain rather resembles that of a tent (B).

The coding metaphor tries to match the causal structure of dominoes to the causal structure of a tent (Fig. 10), but they are not congruent. In fact, the causal structure of the brain is more complex than that of a tent, because in addition to the coupling of neurons, the brain itself is coupled to its environment, i.e., there is circular and not linear causality (Fig. 9A). As Dewey (1896) pointed out more than a century ago: *“the motor response determines the stimulus, just as truly as sensory stimulus determines the movement.”*. Many other authors in biology, psychology, philosophy and robotics have argued that perception is not a one-way process but an interaction with the environment (Powers, 1973; Gibson, 1986; Brooks, 1991; O’Regan and Noë, 2001; Ahissar and Assa, 2016). This makes the notion of stimulus questionable; but the neural coding metaphor implies that neural activity is responses to stimuli, not exploration of the environment. How does the linear causal model accounts for a rat exploring a maze with its whiskers? Where does spontaneous activity fit in the coding model? As a theory of cognition, the neural coding metaphor seems to embrace the most basic form of behaviorism^10^.

## 5. Conclusion

### 5.1. Summary

When I say that the heart is a pump, I propose a function for the heart (to circulate blood) and mechanisms by which blood is circulated; I propose specific ways in which elements of the heart interact by identification with elements of a pump. In effect, the pump is a model of the heart. A metaphor is not just words arbitrarily chosen to designate an object: it is a model of the object (Lakoff and Johnson, 1980), and as such it deserves scrutiny as any other model in science. Is coding a good model of brain function?

There are three aspects of the coding metaphor: correspondence, representation and causality. Technical results are based on the first aspect, but their interpretation and claimed significance draw on the two other aspects which are not subject to the same scrutiny. Many neural coding theories rely on the idea that the brain manipulates neural representations of stimulus properties, as if the variable of a neural code were a processor register that the brain can store, retrieve and combine arbitrarily, while knowing what the variable refers to. But what is the evidence that such neural representations exist, and what is the evidence that the brain can manipulate spikes in this way?

Technically, it is found that the activity of many neurons varies with stimulus parameter, but also with sensory, behavioral and cognitive context; neurons are also active in the absence of any particular stimulus. A tight correspondence between stimulus property and neural activity only exists within a highly constrained experimental situation. Thus, neural codes have much less representational power than generally claimed or implied. Behavioral significance is only obtained by making an implicit *“linking proposition”* (Teller, 1984) that relates coding variables and behavior, which takes the form of a “decoder”. The decoder, often an “ideal observer”, is a hypothetical abstract construct whose biological basis is unspecified and whose existence is unquestioned, even though the decoder must incorporate key contextual aspects, including methodological details of the experiment which defines the coding variables. Critically, the contextual dependence of neural codes cannot be solved by incorporating contextual variables in a broader neural code, because context is precisely what defines the variables. A perceptual scene cannot be fully defined as a vector of properties; properties of what?

The notion of information implied by the coding metaphor is inappropriate to understand perception and behavior, because it is information by reference to external symbols. A more appropriate notion is information as organization (Thompson, 1968), namely relations between sensory signals and actions, forming a structured internal model. The relation between such structured models of the world and neural activity is unclear, but what is clear is that no neural coding theory proposed so far seems adequate even in principle.

Ultimately, the neural coding metaphor is a way to think about the brain that is disconnected from its causal structure. The brain is a dynamical system coupled to the environment, and is itself composed of coupled dynamical systems (neurons), whose interaction is mediated by spikes, which are timed events. The dualistic structure of the metaphor cuts through this organization and decides that one part of the brain can be understood independently of the way it interacts with the rest of the brain, and independently of the way the brain interacts with the world. More fundamentally, a causal role is attributed to coding variables, but this is incoherent because coding variables are extended measurements of activity linked to the temporality of experiments; they are not causal variables of the underlying dynamical system. In conclusion, the causal structure of neural coding metaphor is incongruent with the causal structure of the brain. If neural codes have no causal power, then they cannot form the basis of a theory of brain function.

### 5.2. What else, if not coding?

Since the coding metaphor is so ingrained in neuroscience, how could it be possible to abandon it? What could it be replaced with?

First of all, that there is no simple substitute for neural coding does not make it a viable option. The neural coding metaphor is attractive partly because it resonates with Cartesian philosophy, as pointed out by Cisek (1999), and partly because it seems to fit with the computational view of the mind, the idea that cognition is the manipulation of symbols that represent properties of objects in the world. But the symbols provided by neural codes are not context-free, they are unstructured and they have no causal powers. They do not have the quality required by the computational view. Thus, the appeal of the neural coding metaphor is illusory. Even if it were possible to map brain activity to computational descriptions, neural codes would not provide the adequate mapping.

The alternative route is to embrace the causal structure of the brain and to adopt a systemic approach. The brain is a system; or more accurately the brain, body and environment are a system. This approach is precisely what the coding metaphor forbids, since it cuts through the system and uncouples its different components. Since it is a dynamical system, this view is related to the dynamical view of cognition (Gelder, 1998). But the specific point here is not so much that cognition is dynamic, but rather that its neural basis is a dynamical system and must be understood as such. It is a special kind of dynamical system, in that it is composed of units (neurons) which are also dynamical systems. The causal role of spikes in this system is to mediate coupling between these dynamic units. They are transient events that are better understood as actions than as representations (a representation is not an event).

In terms of neural modeling, this requires considering sensorimotor systems. Paradoxically, it is customary in systems neuroscience to model perceptual abilities by considering only the corresponding sensory areas. We speak for example of the visual system as a set of anatomical structures from the eye to the visual cortex. But the visual system defined in this way is not actually a system, if it is disconnected from the elements without which it cannot have any function. It follows that models of perceptual systems are in effect not biological models, but chimaeras obtained by attaching a neural model of a sensory area to an abstract construct (“decoder”) that maps the activity of neurons to descriptors of behavior, and often to an even more problematic abstract construct (“encoder”) that maps stimulus parameters to model inputs. This methodology embraces both behaviorism (neural activity is only responses to stimuli) and dualism (something else makes sense of neural activity). Instead, I suggest developing models of the full sensorimotor loop, “models that behave” (Gomez-Marin, 2017). For example, instead of looking for neural codes of sound location, one could look for neural models of auditory orientation reflexes. Measurements of neural activity in stimulus-response experiments can be used to constrain and test such models, but they do not need to be the output of the model, nor do they need to be a causal variable in the model. To be clear, the issue is not about the amount of detail that needs to be incorporated. Models can be simplified or idealized, as any model needs to be. The issue is to respect the causal structure of brain and behavior, and to see neural activity as what it really is, activity. Action potentials are potentials that produce actions, they are not hieroglyphs to be deciphered.

## Footnotes

1 A strong correlation (or mutual information) between wavelength and current observed in the first case (Fig. 2B) may transfer to a negligible correlation in the second one (Fig. 2C) (Brette, 2010).

2 A similar number of neurons are also tuned to small ITDs, especially in larger mammals such as cats (Goodman et al., 2013, Fig. 1a).

3 Suppose for example that we observe the activity A of a neuron, whose firing rate varies with parameter X as follows: A = X + Z, where Z is an uncontrolled variable. If Z has large variance, A might be hardly correlated with X. But if we simultaneously observe B = Z, then we can recover X exactly (B-A), even though B is not correlated at all with X. There is no direct relation between parameter sensitivity assessed by a tuning curve and information in a broader context.

4 For example, Jazayeri and Movshon (2006) argue that the representation of the probability distribution of stimulus property (rather than just the most likely property), allows the same code to perform different tasks, and comment: “In contrast, previous models of sensory decoding were for the most part designed to account for a particular task.”. But the new proposition still requires a specific decoder for each task.

5 Not to be confused with predictive information, which is the mutual (Shannon) information between the past and future of a signal (Bialek et al., 2001; Palmer et al., 2015).

6 This proposition remains controversial, because it is unclear whether synchronous firing across distant brain areas has causal powers (Merker, 2013) (see next part).

7 Nonetheless, the resemblance with Cartesian dualism is hard to miss. Indeed, Cisek (1999) argues that this dualistic structure has been inherited from Cartesian dualism, specifically that computationalism has replaced the non-physical mind by a mechanistic cognition, while keeping the architecture unchanged (perception-cognition-action).

8 Except if the law of large numbers is used. This requires a number of assumptions, see Brette (2015).

9 A notable exception being, of course, a computer executing an algorithm.

10 A variation that Gomez-Marin calls “neuralism” (Gomez-Marin, 2017.

